# Emergent Diversity and Persistent Turnover in Evolving Microbial Cross-Feeding Networks

**DOI:** 10.1101/2021.12.17.472886

**Authors:** Leonhard Lücken, Sinikka T. Lennartz, Jule Froehlich, Bernd Blasius

**Affiliations:** Institute for Chemistry and Biology of the Marine Environment (ICBM), Carl von Ossietzky University of Oldenburg, Oldenburg, Germany; Department of Earth, Atmospheric and Planetary Sciences, Massachusetts Institute of Technology, Cambridge, MA, USA; Helmholtz Institute for Functional Marine Biodiversity (HIFMB), Carl von Ossietzky University Oldenburg, Oldenburg, Germany

**Keywords:** microbial diversity, cross-feeding, metabolite exchange, consumer-resource systems, community dynamics, trophic networks, evolving networks

## Abstract

A distinguishing feature of many ecological networks in the microbial realm is the diversity of substrates that could potentially serve as energy sources for microbial consumers. The microorganisms are themselves the agents of compound diversification via metabolite excretion or overflow metabolism. It has been suggested that the emerging richness of different substrates is an important condition for the immense biological diversity in microbial ecosystems. In this work, we study how complex cross-feeding networks (CFN) of microbial species may develop from a simple initial community given some elemental evolutionary mechanisms of resource-dependent speciation and extinctions using a network flow model. We report results of several numerical experiments and report an in-depth analysis of the evolutionary dynamics. We find that even in stable environments, the system is subject to persisting turnover, indicating an ongoing co-evolution. Further, we compare the impact of different parameters, such as the ratio of mineralization, as well as the metabolic versatility and variability on the evolving community structure. The results imply that high microbial and molecular diversity is an emergent property of evolution in cross-feeding networks, which affects transformation and accumulation of substrates in natural systems, such as soils and oceans, with potential relevance to biotechnological applications.

## 1 Introduction

Microorganisms have successfully inhabited both aquatic and terrestrial ecosystems on earth for millions of years, and continuously evolved to an enormous diversity [1-5]. This immense ecological success is related to the observation that microorganisms are able to modify their abiotic environment through the production of organic compounds. Thereby, due to their metabolic activity, microbes act as ecosystem engineers and over the millennia have been able to completely transform and alter the chemical environment on the planetary surface, with drastic consequences for conditions of life and biogeochemical cycles [6]. In the marine environment, for example, microbial transformation of organic compounds is considered to have played an important role in the production of refractory matter, creating a pool of dissolved organic material that resists biodegradation over thousands of years and has accumulated in the oceans to a current reservoir of 700 Petagram carbon - with strong implications for the global climate system [7-9].

Based on their enormous diversity, and mediated by the exchange of metabolites, microbial communities are forming incredibly complex networks of ecological interactions [10, 11]. In many cases, a substance that a species may or must use for its growth, is excreted by another species as a byproduct, yielding syntrophic or cross-feeding relationships [12-15]. Recent evidence suggests that such cross-feeding interactions should be a generic feature of microbial ecology 16] and it is believed to be partly responsible for difficulties of cultivating a large portion of microbial species occurring in natural habitats [17]. The full importance of cross-feeding for microbial biodiversity is, however, still a matter of debate [18].

One other defining characteristic of microbial communities is their potential for fast evolution. Due to fast turnover times and the possibility of lateral gene transfer [19] new mutations occur frequently even on laboratory time scales [20]. The potential for rapid evolution of metabolic cross-feeding interactions was convincingly demonstrated in a series of long-term experimental studies with microbial populations [21-23]. This was also confirmed in model studies, showing that cross-feeding should naturally emerge if waste products or less valuable compounds are generated during the metabolization of a primary resource [24, 25]. Additional evidence for naturally occurring rapid evolution of new metabolic pathways comes from observations of microbial degradation of organic pollutants that have been introduced by humans only decades ago [26]. For instance, in the marine environment, the increased introduction of plastic waste created a new ecological niche, the ‘plastisphere’, which serves as colonization habitat for diverse microbial communities, with some of them already having developed enzymatic pathways that can hydrolyze and degrade certain plastic polymers [27-30].

In diverse microbial communities, such evolutionary inventions will scale up, giving rise to ongoing alteration of metabolic transformations. This dynamic is further complicated by the fact that an altered chemical environment will affect the fitness of individual organisms, leading to extinctions of microbes that have lost essential substrates, while the production of new metabolites provides niches for either speciation or invasion of new microbial consumers - again altering the chemical environment and yielding successive extinction and invasion events. In this way, microbial and molecular diversity mutually sustain each other, in the sense that microorganisms diversify compounds by metabolite excretion and overflow metabolism, and a diverse set of metabolites offers new niches for evolving microbes. In order to gain an understanding of the resulting complex co-evolutionary dynamics, theoretic approaches have proven to be a powerful method.

One of the most influential conceptual models of large-scale co-evolution in this context is a model introduced by Bak and Sneppen [31], which directly assigns a new random value for the fitness of the least fit population and all interacting populations, mimicking the replacement of the former and the alteration of its interactions. Later, Solé and Manrubia proposed a similarly simple model of an ecological network, considering a more continuous drift of properties and ancestral relations between populations [32, 33]. Just as the Bak-Sneppen model, it assumes a fixed number of species. Nunes Amaral and Meyer [34] then considered foodweb models and allowed their size and depth to vary, driven by the processes of extinction and speciation.

These conceptual models, based on simple statistical rules, focused to determine the distribution of extinction cascade sizes. They did not, however, allow to simulate the time dependence of consumer and resource densities. This is possible in another class of models that extend classical consumer-resource models [35]. Besides including resource consumption, models of this class also consider the exudation of compounds by different consumer populations. In turn, these products provide resources available to other populations, which possess metabolic capabilities for consumption. We denote the resulting bipartite network of consumers and resources, in which resource-to-consumer links represent consumption and consumer-to-resource links production, as cross-feeding networks (CFN).

The analysis of CFN models received an increasing attention in the past few years [16, 36-42]. The usual approach for assembling a model community in CFN models is the creation of a meta-population pool, holding the full entirety of existing species. In this step, structural assumptions regarding feasible substrate transformations and consumer traits take effect. Here, energetic and stoichiometric relationships between different compounds or correlations between different metabolic capacities may be considered. This step is then followed by placing a subset of species in a given environment and applying environmental filtering to determine a resulting CFN, which then could represent observed patterns in natural systems [16, 39, 40, 43].

These studies, however, did not consider evolutionary dynamics. In turn, most of the experimental and modeling studies concerned with the evolutionary emergence of cross-feeding were restricted to situations where only two consumers species interact[24, 25, 44]. An exception is the study of Goyal and Maslov [37], who modeled how a trophic structure involving many, mostly hierarchically organized, consumers can arise mediated from metabolite leakage. In their model, a one-to-one relationship is assumed for consumer species and resources, which restricts the degree of reciprocate interaction in the generated networks. In the progress of the evolution, larger changes become rarer and rarer in the model, as the system develops towards a saturated or optimized state. Besides the work of Goyal and Maslov 37], the effect of evolutionary processes on complex CFNs has – according to our state of knowledge – not been studied in detail yet.

Given the commonness of CFNs in natural systems [14], understanding the evolutionary dynamics of CFNs and separating between internal and external variability is crucial. Is the continuous saturation observed in the model of Goyal and Maslov [37] an inherent property of the evolution of CFNs? Or is it a result of the models simplifications, such as the limitation of the trophic dependencies of a consumer to a single resource? In natural environments, microbial consumers thrive on a variety of different substrates, implying a higher in-degree of the consumer nodes in a network representation than assumed in [37].

In our work, we consider the dynamics of an evolutionary CFN model allowing higher diversities of the consumed and released resources. Without considering the underlying biochemical foundations, we study the emerging network structures and the time course of their evolution. Our protocol is capable of describing the appearance of novel resources and the corresponding rise of novel metabolic capabilities to exploit these. We report results of several numerical experiments and provide an in-depth analysis of the evolutionary dynamics. Further, we compare the impact of different parameters, such as the assumed metabolic efficiency, versatility, and variability on the evolving community structure. Our findings show that a complex ecological system, which is subject to persisting turnover even in stable environments, can arise from simple assumptions.

The paper is organized as follows. In Sec. 2, we introduce the model, which comprises a formalism for the calculation of stationary flows in a CFN and a protocol describing the evolutionary development of the network. In Section 3, we provide theoretical results on the maximal system capacity and a numerical analysis of the evolutionary process. In particular, we determine the parameter dependence of stationary flow patterns, as well as structural properties and biodiversity measures of the evolving system. In the final Section 4, we discuss the relation of our model and its displayed characteristics to previous experimental and modeling studies.

## 2 Materials and Methods

### 2.1 Stationary Flow in CFNs

Cross-feeding networks are directed bipartite networks composed of two different types of nodes: consumers and resources, cf. Fig. 1. Thus, two different link types and, correspondingly, two different types of resource flow exist in the network. Links directed from resources to consumers indicate the uptake of the corresponding resource by the corresponding consumer, whereas a link from a consumer to a resource describes the release of the resource by the consumer. Deliberately, we do not specify a physical unit for the flow, since the model is generic with respect to different choices thereof. For instance, as a physiological equivalent of the network flow, one may consider moles of carbon transferred from one node to the other, or being exported to inorganic or accumulating reservoirs. Although we employ this notion of mass flow throughout this work, one could likewise consider the energy bound in the chemical composition of compounds to constitute the unit of measurement [37]. We refer to the flow from resources to consumers as “uptake” and to the flow from consumers to resources as “release”. More formally, in a given CFN we denote the set of all resources by ℳ and the set of all consumers by *𝒩*. The uptake rate of resource *j* ∈ *ℳ* to consumer *i* ∈ *𝒩* is then denoted by *J*_*j*→*i*_ and the release rate of resource *j* from consumer *i* as *F*_*i*→*j*_, see Fig. 1 B. The total uptake rate at consumer *i* is the sum

**Figure 1:**
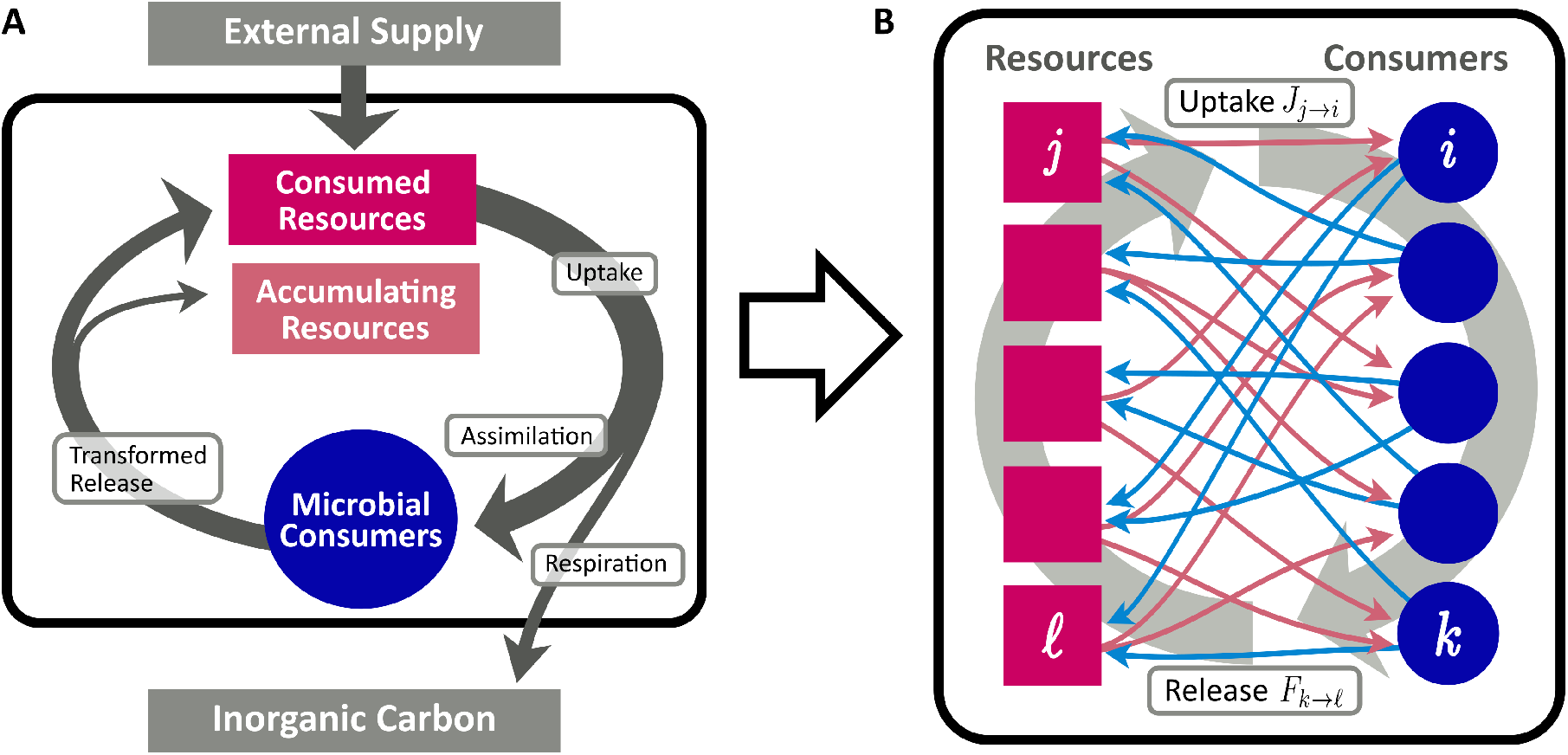
Conceptional diagram of the CFN model, describing the flow of organic matter (arrows) between pools of resources (red rectangles) and microbial consumers (blue circles). Panel A: Aggregated flows between different pools. Organic matter is supplied from an external source to the resource pool. A fraction *η* of consumed resources is assimilated for microbial growth, while the remainder is lost from the system and remineralized to inorganic carbon. Assimilated resources are eventually recycled and enter the resource pool in transformed molecular composition. Panel B: Fine grained scheme of the bipartite network of assimilation red arrows, cf. Eq. (4)] and release blue arrows, cf. Eq. (6)].

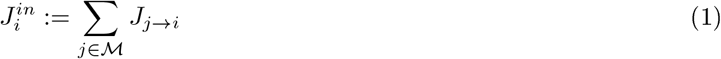

of all individual uptake rates.

Besides the release originating in consumers, we allow external supply flows *s*_*j*_ at each resource, such that the total inflow rate at resource *j* is

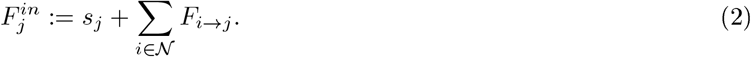

Our modeling approach assumes a given set of consumers at each moment in evolutionary time. Consumers have specific metabolic profiles, which determine their interaction with the present resources. Each consumer *i* is equipped with a vector of resource affinities *u*_*i,j*_, *j* ∈ *ℳ* and a vector of release proportions *ϱ*_*j,i*_, *j* ∈ *ℳ*, with ∑_*j*∈ ℳ_ ϱ_*j,i*_ = 1.

The share *σ*_*i,j*_ ∈ [0, 1] of resource *j* acquired by the *i*-th population is considered proportional to its affinity for that resource under the requirement that ∑ _*i*∈𝒩_ *σ*_*i,j*_ = 1. This gives

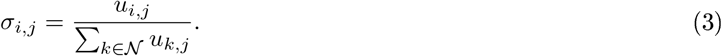

If no consumer exists for a specific substrate at a given time, i.e., *u*_*i,j*_ ≡ 0 for all *i* ∈ *𝒩*, we set *σ*_*i,j*_ to zero for all *i*. In general, a certain fraction *γ*_*i,j*_ ∈ (0, 1) of a consumed resource is remineralized during respiration, while a fraction *η*_*i,j*_ = 1 *− γ*_*i,j*_ maintains an organic form, which subsequently re-enters into the resource cycle. For simplicity, we assume homogeneous recycling fractions *η*_*i,j*_ ≡ *η*, and thus obtain the resulting uptake flow *J*_*i,j*_ from resource *j* to consumer *i* as

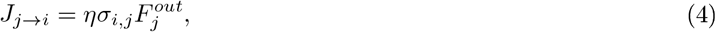

where 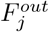 is the total consumption rate of resource *j*. In the following, we consider stationary flows, where for each node, the inflow and the outflow balance, such that 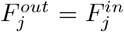, cf. (2). Therefore, we omit the superscripts in the following and simply write *F*_*j*_, resp. *J*_*i*_, for the stationary flows through resources, resp. consumers. Using (1) and (4), the flow at consumer *i* is

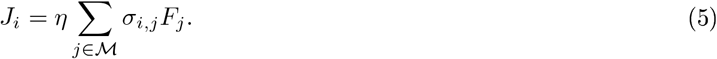

The composition of release by consumer *i* is assumed to follow the release proportions *ϱ*_*j,i*_, *j* ∈ *ℳ*. We assume that the flow *F*_*i*→*j*_ is proportional to the stationary flow *J*_*i*_ through consumer *i* multiplied by the proportion*ϱ* _*j,i*_ released to resource *j*, i.e.,

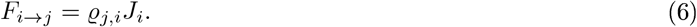

Here, we assume the products to be independent of the composition of the consumption. Combining (2), (5), and (6), we find

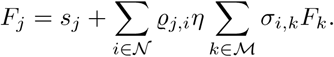

Or in vector, resp. matrix, notation:

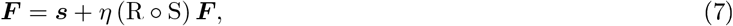

with vectors *F* = (*F*_*j*_)_*j*∈_*ℳ, s* = (*s*_*j*_)_*j*∈_ℳ, and matrices R = (ϱ _*j,i*_)_*j,i*∈_ℳ _×𝒩_ and S = (*σ*_*i,j*_)_*i,j*∈N ×M_. The concatenation T ≔ R ° S of uptake and release partitioning represents the resource transformation matrix in the projection of the bipartite system to the unipartite resource space.

Solving for *F* yields

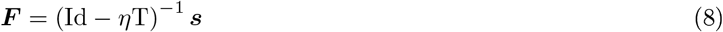

Since the mapping T represents merely a redistribution of the flow across the resource network, it will never increase the total flow, i.e.,

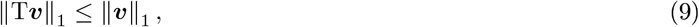

for all *v* ≥ 0, where ‖*v*‖_1_ = ∑_*j*_ |*v*_*j*_| denotes the *L*^1^-norm of *v*. Because 0 *< η <* 1 holds, the inverse of Id – *η*T exists and we may rewrite (8) as

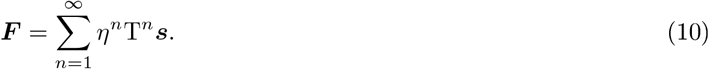

Thus, the internally generated network flow is obtained by the iterative transport of the supply flow through the network. Note the formal relationship of Eq. (7) to the definition of the weighted Katz centrality [45, 46] associated with the resource transformation matrix T. This has a simple interpretation: the stationary resource flow through consumer *i* is proportional to the weighted sum of flows through the network neighbors of *i* plus the input flow from the external source.

### 2.2 Evolutionary Generation of CFNs

While in the previous section we determined the stationary resource flow for a given CFN, we now define the rules that lead to evolutionary changes in the CFN. In our protocol, the change of the CFN within one evolutionary time step is determined by two stages: (1) The randomized generation of a new consumer species with an ancestor in the resident community, and (2) the calculation of the impact of its introduction on the CFN, including eventual extinctions and a modification of the network flow pattern.

The procedure assumes that at the evolutionary time scale, where adaptations of the metabolic capabilities and introductions of new consumer species occur, is separated from the timescale of ecological processes that lead to an equilibration of the network flow. Furthermore, we avoid the calculation of population densities and resource concentrations by describing the system exclusively by the network flow.

In the first stage, a new consumer species *i* is introduced into the community. It is assigned an ancestor *a* in the resident community, which is chosen at random, but with a probability weight proportional to *J*_*a*_. The descendant *k*’s characteristics are then altered on the basis of the ancestor’s. More precisely, given the affinities *u*_*a,j*_ and release fractions *ϱ* _*j,a*_ of the ancestor, the corresponding characteristics of the descendant are set proportional to

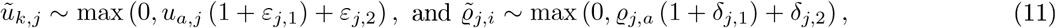

where *ε*_*j*,1_ and *δ*_*j*,1_ are independently, uniformly distributed in (*−α, α*) and small equalizing drifts *ε*_*j*,2_ ≥ 0 and *δ*_*j*,2_ ≥ 0 act upon the present values, i.e.

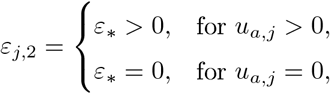

and

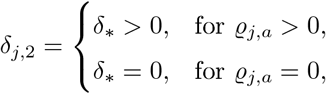

with random variables *ε*_∗_ and *δ*_∗_ drawn from a uniform distribution 𝒰 (0, *β*) with drift intensity *β*. The definite values for the characteristics of the new consumer are generated by drawing them according to (11) and normalizing the resulting totals as

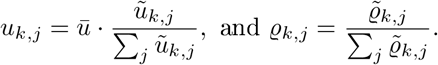

Limiting the total affinity may be considered to represent the limited investment each consumer may place on transporter enzymes and other metabolic machinery associated with the utilized resources [39, 47].

A second possible substep of the first stage, is executed with a probability *p*_*u*_. Here, we allow for a qualitative change of the new consumer’s metabolism by substituting one utilized resource *j*, i.e. *u*_*i,j*_ *>* 0, for another *j*^′^ with *u*_*i,j′*_ = 0. That is, after the substitution resource *j*_*′*_ takes the affinity formerly assigned to *j*, resulting in *u*_*i,j′*_ = 0 and *u*_*i,j*_ *>* 0. In contrast to the gradual changes of affinities and release fractions in (11), this qualitative adjustment of metabolism mimics a mutation allowing the species to exploit a novel resource. Similarly, with a probability *p*_*Q*_, we alter the release configuration to substitute an old product for a new one drawn at random from a global resource pool 𝒢, representing the entirety of possible organic compounds produced in metabolic processes. This step may introduce resources into the system, which were not present before. As a result, a potential niche may be created for future species to settle. Note, that this procedure fixates the number of positive affinities *n*_*u*_ = # {*u*_*i,j*_ *>* 0} and of *n*_*ϱ*_ = # { ϱ_*i,j*_ *>* 0} for all consumers of an ancestral lineage, which we refer to as uptake diversity and release diversity hereafter.

In the second stage of an evolutionary time step, we add the newly generated consumer species to the resident community *𝒩* to obtain an extended community *𝒩* _*′*_. In order to measure the fitness of the present consumers species, we calculate the stationary uptake flows *J* = (*J*_*i*_)_*i*∈𝒩 *′*_ using (5) and (10). We assume that a species, whose uptake flow falls below a threshold *µ >* 0, faces an acute risk of extinction. That is, if one or several species exist, such that

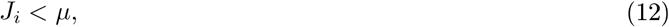

the species with lowest *J*_*i*_ below *µ* is considered extinct and removed from *𝒩* _*′*_, implying an alteration of the flow pattern. The proposed protocol then successively updates the stationary flow and removes the consumer with lowest flow. This procedure is iterated until all uptake flows are larger than *µ*. The resulting CFN is considered to represent a stable community, which now serves as the initial point for the subsequent evolution step. Accordingly, the community evolves in discrete time steps, yielding a sequence of stable CFNs 𝒞_0_, 𝒞_1_, 𝒞_2_, …, where each step introduces a new consumer species and evaluates whether it can successfully establish.

Figure 2 illustrates an evolving CFN at different steps in evolutionary time. It is obtained from the stepwise evolution starting at a single species at *t* = 1 up to *N* = 95 species at *t* = 400. The evolutionary algorithm operates with parameters *n*_*u*_ = *n*_*ϱ*_ = 3 and *p*_*u*_ = *p*_*ϱ*_ = 0.2, and a single resource is externally supplied. Note that the single founding consumer species (blue dot in the leftmost graph) is also capable to consume *n*_*u*_ = 3 different resources, although only one supplied resource is present and depicted, initially. In the subsequent steps, additional consumers are generated. As metabolic novelties evolve, new compounds (red squares) are released and metabolized by the growing community. Clusters of consumer nodes can be observed, which share similar metabolic properties. In Sec. 3.4, we study the clustering in more depth.

**Figure 2:**
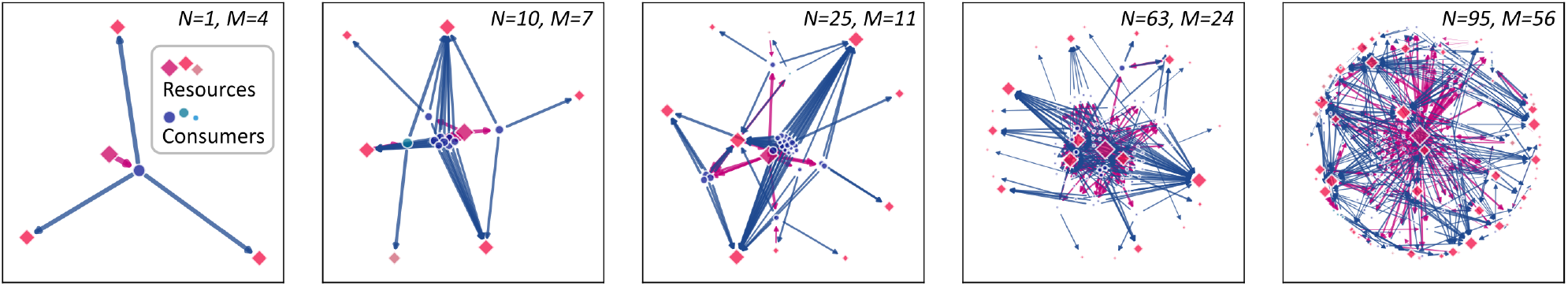
A growing cross-feeding network. Starting from a founding consumer (leftmost panel), new consumers (blue dots) are added sequentially leading to an increasing community size *N*. As novel metabolic capacities evolve, the number *M* of different resources (red squares) increases. The symbol sizes are scaled proportional to the stationary flow passing through the corresponding network node. Uptake links are drawn as magenta arrows, release links as blue.

## 3 Results

### 3.1 Theoretical system capacity and total flow

Due to respiration losses and the finite external supply rate, the number of consumers, which can be sustained in the CFN model, is finite. To determine an upper bound for the number of consumers, let us consider the total flow rate 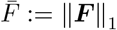 for a given total supply rate 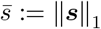. By (10) we have

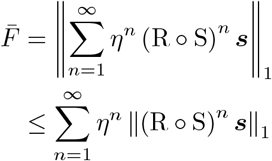

Then, (9) implies

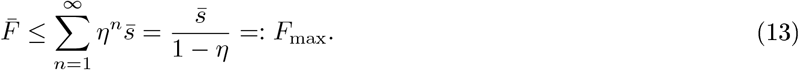

Note that by definition ‖ R*w* ‖_1_ = ‖ *w* ‖ _1_ for all *w* ≥ 0, because all flow assimilated into consumer pools is released into resource pools again. Thus, the inequality (13) may become strict, if ‖ S*v*‖ _1_ *<* ‖ *v* ‖ _1_. This would mean that not all of the resource flow is consumed. A resource *j* without consumers but positive inflow *F*_*j*_ *>* 0 will eventually accumulate and be involved in other processes. This may either be an external degradation, which we do not consider here, or an integration into the recycling process by the arrival of a consumer, which is able to utilize the resource *j*. These accumulating resources are treated as sinks in the model. The same holds for flow entering the inorganic pool through respiration, cf. Fig. 1. Denoting the set of accumulating resources by *𝒜*, the total accumulation flow is then

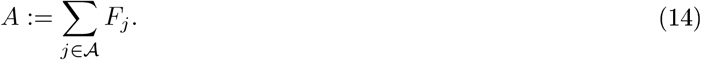

If all resources were consumed, an additional flow of *ηA* would enter the system via the consumption of the accumulating resources and generate an amount

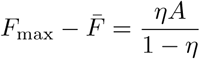

of additional flow until its complete respiration. Taking this into account, we can state more precisely:

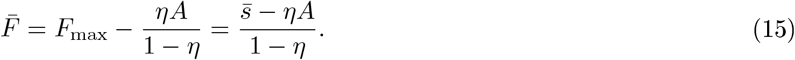

By definition, the total uptake flow 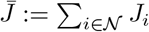 is given by

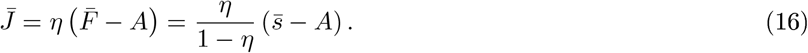

Because a consumer population is only considered persistent if (12) holds, the total number of consumers cannot exceed 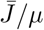. That is,

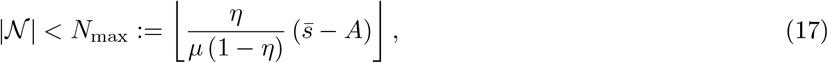

where ⌊ *x* ⌋ denotes the integer part of *x* ∈ ℝ_+_. To obtain an estimate independent of the network structure, one may formally set *A* = 0 in (17).

### 3.2 Simulation results

In the following, we consider the simulation of the evolution of a CFN according to the protocol defined in Section 2 with parameters as defined in the caption of Fig. 3. We assume that the external supply enters the pool of resource *j* = 0 at a fixed rate *s*_0_ ≡ 1.0, mimicking a static environment, which allows us to study the internal dynamics of the CFN. Panels A-F of that figure show the evolution of several measures for an evolving CFN initialized with a random founding species, which has positive affinities for the supplied substrate (and two other substrates not present at the beginning of the simulation). Its release diversity is set to *n*_*ϱ*_ = 3 and is inherited to all its descendants. The time series are split up into a magnified view of the initial phase and a larger, later interval when the system exhibits statistically stationary fluctuations.

**Figure 3:**
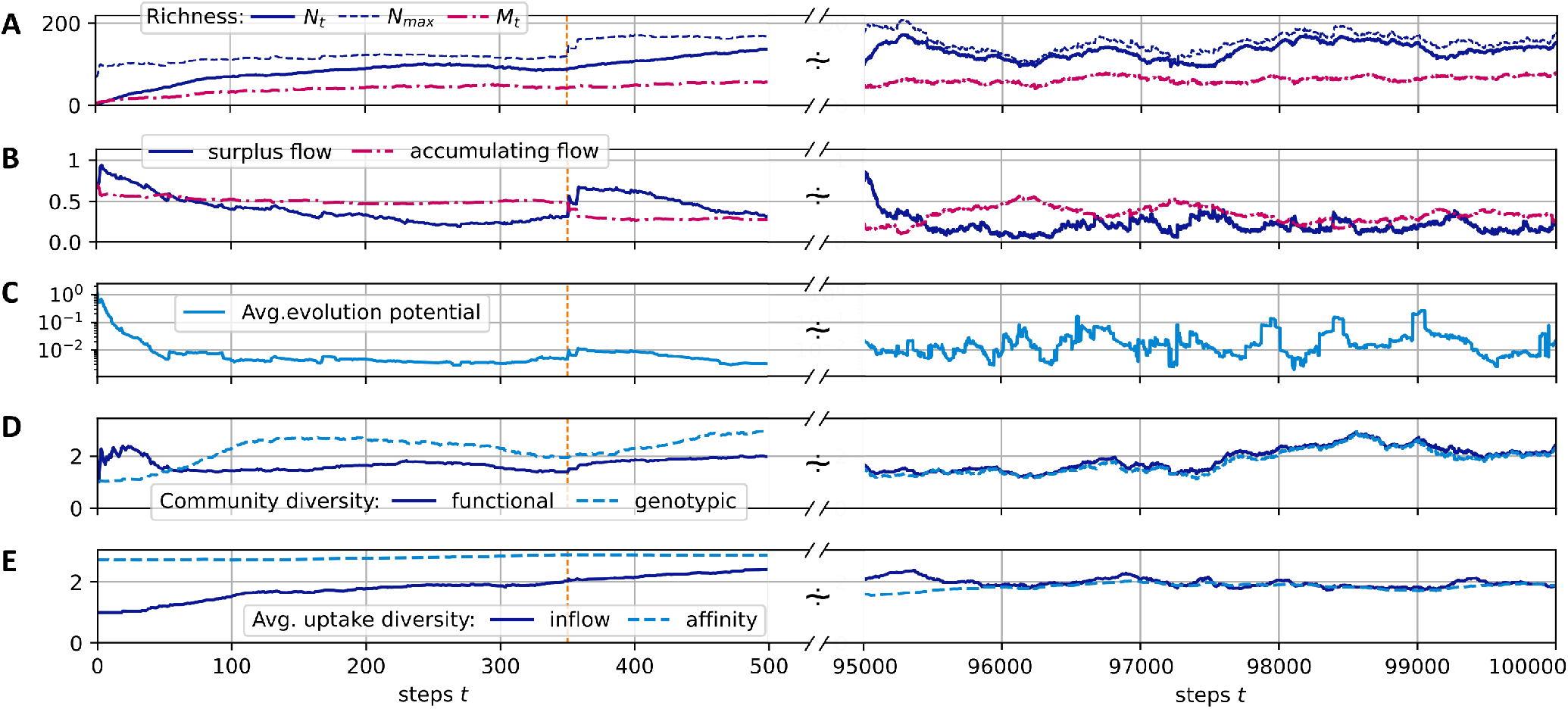
A-E: Different community measures over the course of a simulated evolution. The initial time interval of steps 0 to 500 is magnified, with a dashed orange vertical line indicating step 350. An interval [500, 9.5 × 10^4^] is skipped and the interval [9.5, 10] × 10^4^ is displayed at the right side. See the main text for the definition of the shown quantities. System parameters are *n*_*u*_ = *n*_*ϱ*_ = 3, *η* = 0.7, *µ* = 0.01, *p*_*u*_ = *p*_*ϱ*_ = 0.2, *α* = 0.2, *δ*_∗_ ∼ 𝒰 (0, *α/n*_*ϱ*_), *ε*_∗_ 𝒰 (0, *α/n*_*u*_), 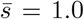 (with only resource being supplied), and # 𝒢= 250. A single founding consumer was inserted at initialization.

#### Richness

Panel A of Fig. 3 illustrates the development of the consumer and substrate richness, *N*_*t*_ = |ℳ _*t*_| (blue, solid curve) and *M*_*t*_ = |ℳ _*t*_| (red, dash-dotted curve), where the subscript *t* references the state at step *t*. The dashed, blue curve shows the upper bound *N*_max_ to the total population size, see (17). For the consumer and resource richness, we can observe an initial increase, which slows down until stationary fluctuations around *N*_*t*_ ≈ 125 and *M*_*t*_ ≈ 65 are reached. After the initial filling of the system, the consumer and resource richness exhibit bounded fluctuations around those values.

#### Network flow

As shown in Panel A, the realized richness remains below the upper limit *N*_max_. This implies that either the uptake rates of some consumers significantly exceed the subsistence flow, i.e., *J*_*i*_ ≥ *µ*, or some of the network flow leaks out of the system into resources, which are not consumed, i.e., *A >* 0. Panel B shows that both is the case, in general. Neither does the accumulation flow *A* drop to zero, nor does the “surplus flow” ∑_*i*∈𝒩_ (*J*_*i*_ *− µ*). At step 350 (indicated by the orange vertical line), a redirection of previously accumulating resources back into the system results in a greatly increased surplus flow, as can be seen by the “jumps” in both quantities in that step. This corresponds to the entrance of a new consumer with the metabolic ability to degrade a resource previously contributing to the accumulating fraction of the flow. Similar events occur regularly also in the evolved states at later times.

#### Evolutionary potential

The fraction of flow into accumulating resources represents a measure for the potential of novel metabolic capabilities to allocate additional resource flow for their bearer. Further, the fraction of surplus flow corresponds to the fraction of flow which can be consumed by additional consumers without inducing a displacement of present consumers. However, if the surplus flow of a specific consumer *i* falls towards zero, it faces the risk of being displaced by another consumer, which has a superior combination of affinities. Any offspring *k*, whose affinities would increase the uptake in comparison to consumer *i* has the potential to do so. Let us assume the substitution of consumer *i* by a similar consumer *k*. Ignoring nonlinear effects and impacts from changing network flow patterns, a first-order estimate can be given for the total uptake flow *J*_*k*_ acquired by consumer *k* in terms of the affinity differences *δ*_*j*_ := *u*_*k,j*_ *− u*_*i,j*_, by

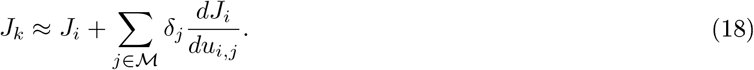

Thus, to measure the potential for increasing the fitness of consumer *i*, one may consider the redistribution of its affinities *u*_*i,j*_ which maximizes (18). Recall, that the admissible variation of the affinities is constrained to preserve the total affinity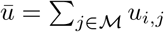. For simplicity, let us assume that the variation keeps the set of uptake capabilities fixed, that is, only *u*_*i,j*_ with *u*_*i,j*_ *>* 0 are allowed to vary. Then, the highest potential increase in the uptake rate by such a redistribution is achieved by shifting the preference from *u*_*i*,min_ with minimal 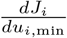 to *u*_*i*,max_ with maximal 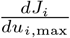. Accordingly, an indicator for the potential evolutionary improvement would be the quantity

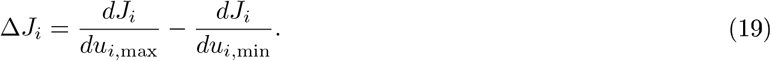

We call Δ*J*_*i*_ the “evolutionary potential” of consumer *i* and we report its average over the whole ensemble *𝒩* in Fig. 3 C.

A straightforward calculation shows that

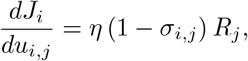

where the value *R*_*j*_ = *F*_*j*_*/ ∑*_*n*_ *u*_*n,j*_.

The convergence of Δ*J*_*i*_ to zero would indicate an evolution towards an optimal, saturated system state where the present ensemble of consumers can hardly be invaded. However, in our simulations, it fluctuates around finite values, indicating an ongoing co-evolution, which constantly creates new niches and results in a persistent community turnover with time. Thus, despite a constant supply flow, the system does not reach an evolutionary stable community.

#### Diversity

Panel D in Fig. 3 shows measures for the genotypic and functional diversity of the consumer community. We calculate these diversity indices based on the pairwise species similarities *z*_*i,k*_, *i, k* ∈ *𝒩*, as proposed by Leinster and Cobbold 48]. To determine the pairwise similarities, we calculate the cosine similarity of either the species’ affinities (*u*_*i,j*_)_*j*∈_ℳ to obtain a genotypic quantity, or of the realized uptake flows (*J*_*j*→*i*_)_*j*∈_ℳ, which gives a measure of functional diversity. Given the pairwise similarities, the ordinarity of a species *i* ∈ *𝒩* is defined as

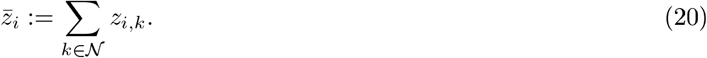

The diversity *D* for the whole community is calculated as the inverse of the average ordinarity, i.e.,

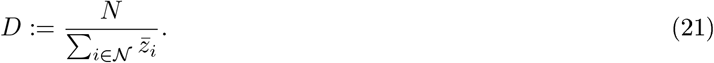

We note that (21) corresponds to the diversity of order *q* = 2 as defined by Leinster and Cobbold [48].

In the simulation run shown in Fig. 3, both diversity measures fluctuate relatively synchronously around an average value of about 2.1. This relatively low value, compared to the average consumer richness of about 125 (see Fig. 3 A), can be explained by the fact that a large cluster of consumers concentrates its efforts on the consumption of the only supplied resource *j* = 0, which has the highest inflow of *F*_0_≳ 1.0, and a few produced resources, which have higher availability. On average two thirds of consumers exhibit an uptake flow, which is composed to more than 1/2 from the supplied resource. See also Fig 4 A, which describes the relationship of evolved affinities and substrate flows, and is discussed below.

**Figure 4:**
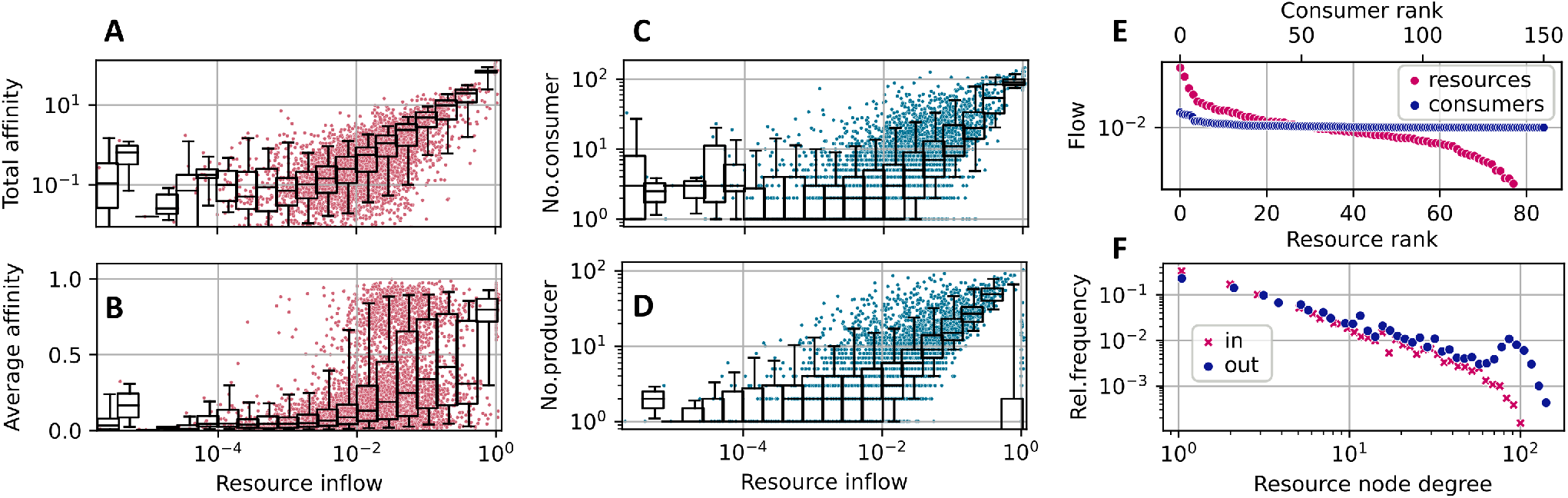
Community and network statistics for the simulation run shown in Fig. 3, taking into account steps *t* ≥ 50000. Panels A-D: Resource flows *F*_*j*_ (*t*), *j* ∈ *ℳ* _*t*_, plotted against: A - the total affinity (22); B - the average affinity *U*_*j*_ (*t*) */C*_*j*_ (*t*), where *C*_*j*_ (*t*) := # {*u*_*i,j*_ *>* 0, *i* ∈ *𝒩*_*t*_} is the number of consumers of resource *j*; C - the number of consumers *C*_*j*_ (*t*); D - the number of producers *P*_*j*_ (*t*) := # { _*j,i*_ *>* 0, *i* ∈ *𝒩*_*t*_}. Panel E: Rank vs. stationary flow diagram for consumers (blue points) and resources (red points) in the community *𝒩*_*t*_ for *t* = 10^5^. Panel F: Resource node logarithmic degree distributions: log(in-degree) density, i.e., #{*P*_*j*_ (*t*) = *k*}, as red crosses; log(out-degree) density, i.e., # {*C*_*j*_ (*t*) = *k*}, as blue dots. All community states *𝒩*_*t*_, *t* ∈ 𝒯 see (23)], are aggregated in E.

Figure 3 E shows the evolution of the average uptake diversity of individual consumers. As a measure of consumer *i*’s position on a range from being a single resource specialist or a generalist, we calculate the effective richness [48]

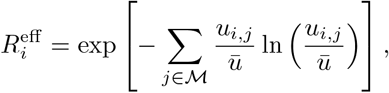

of its relative affinities 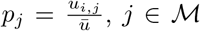, or inflows 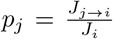, respectively. A value of 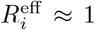 indicates that consumer *i* is a single resource specialist while higher values indicate that the consumer does, or may, thrive on different resources. The maximal possible value for 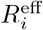 equals the uptake diversity *n*_*u*_ = 3. In the shown simulation, a community average of about 2 establishes, indicating that, besides the fraction of consumers specializing on the supplied resource, others exhibit more generalist properties. See also Fig. 4 B, where we show the distribution of the average affinities of consumers over specific resources.

The relation of the total effort dedicated to the competition for a resource and its availability is illustrated by the statistics shown in Fig 4 A. Here, each red dot shows the relation of the total affinity

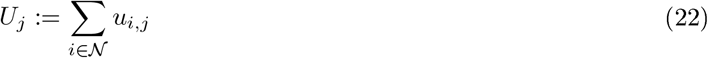

to the stationary flow *F*_*j*_ for a specific resource *j* at a specific time step *t* from a sequence of community states (*𝒩*_*t*_)_*t*∈𝒯_ at 201 equidistant steps

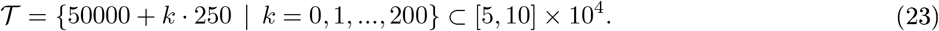

The overlaying boxplot summarizes the distribution of points for the corresponding bins, indicating the median and the 25%-75% quantile range within the box and the 5%-95% range within the whiskers. A linear regression to the double log data restricted to resources with *F*_*j*_ *>* 0.001, yields an exponent ≈ 1.0 for the dependence of total affinity on total flow, indicating a strong linear correspondence. This means that the total effort, i.e., affinity, in the community, dedicated to the uptake of a resource is linearly proportional to its availability. Note, that in a dynamical consumer-resource model with specific uptake rates *u*_*i,j*_, this linear distribution would correspond to a homogeneous distribution of stationary concentrations 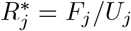.

Similarly, as visible in Fig 4 C and D, the number of consumers increases with the availability of the resource, and the availability of a resource increases with the number of associated producers. For the latter relation, the supplied resource forms an exception, as its inflow is high without necessarily having a large number of producers.

In Figure 4 B, we show the distribution of the average affinities of consumers over specific resources. We observe an interesting pattern of transition from high to low average affinities depending on the resource flows *F*_*j*_. For the supplied resource the average affinity is close to one, indicating specialist consumers. On the other hand, for resources with low inflow, we observe mainly low affinities. In the mid range, though, substrates may bear both, specialists and generalists.

In Panel F we show the distributions of logarithmic in-degrees and out-degrees of the resources node, i.e., of the number of consumers and producers associated with present resources. Here, we observe an approximate power law scaling of relative frequencies with exponents ≈ *−*1.28 (for in-degrees) and ≈ *−*1.08 (for out-degrees) up to degrees ≤ 50. The appearance of higher frequency of out-degrees in the range ≈ [70, 120] corresponds to the persistently high number of consumers associated with the supplied substrate.

Figure 4 E shows an exemplary diagram of rank vs stationary flow in the last community *𝒩*_*t*_ of the simulation run shown in Fig. 3. It is worth noting that the fractions of flow allocated to the different consumers are very evenly distributed with *J*_*i*_ *µ* for all *i* ∈ *𝒩*_*t*_ (the Pilou evenness is ≈ 0.993). This corresponds to a a relatively high degree of saturation in the system, where almost all present consumers are easily driven across the extinction threshold at *J*_*i*_ = *µ* by newly arriving consumers. Figure 5 C shows that this does not lead to extinction cascades of arbitrary sizes, though, as consumers fulfilling (12) are not removed at once. Instead, the removal is performed successively and the stationary flows are updated after each removal. This reduces the competition gradually until *J*_*i*_ ≥ *µ* for all remaining consumers.

**Figure 5:**
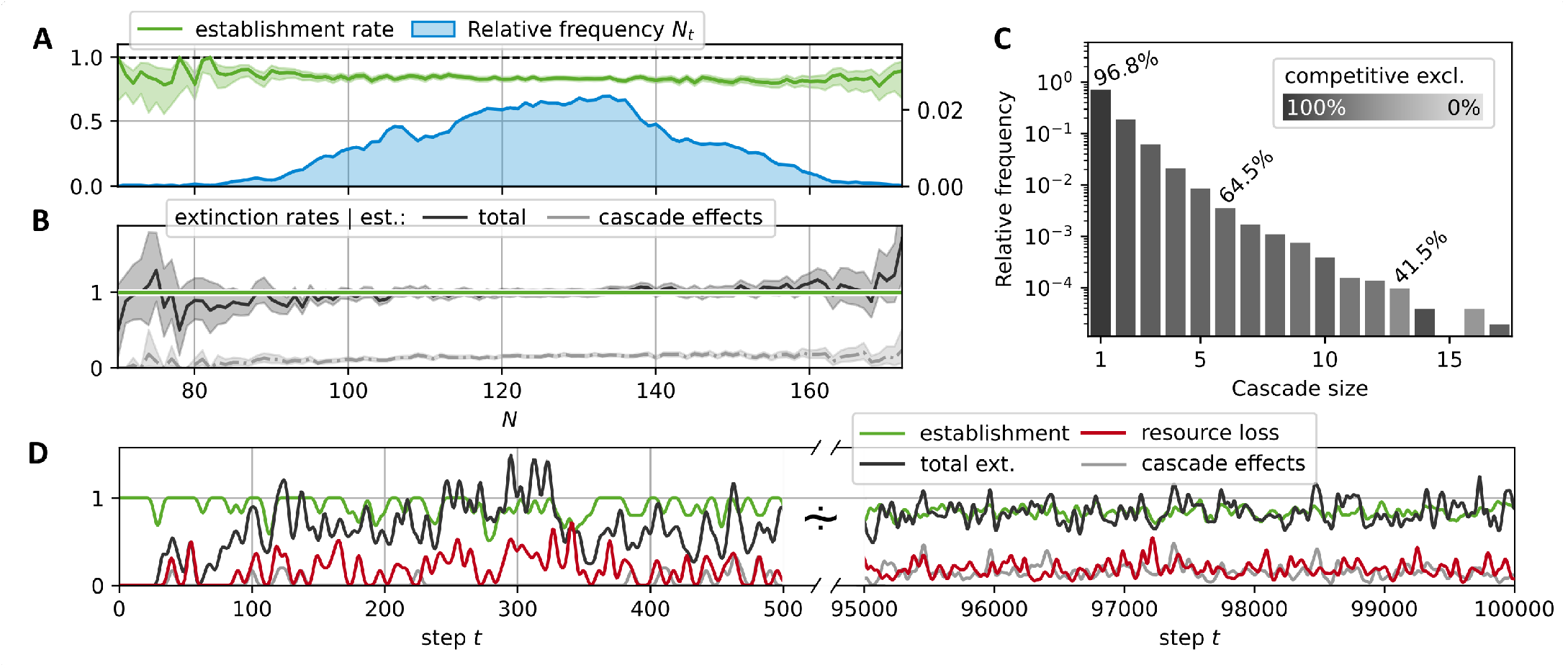
Displacement statistics for the simulation run shown in Fig. 3; A-C take into account steps *t* ≥ 50000. Panel A: Average success rate for the establishment of the new consumer generated in a step *t* given the community size *N*_*t*_ (green curve); observed distribution of *N*_*t*_ (blue curve). Panel B: Average rate of extinctions given a successful establishment of a new species in dependence of the community size *N*_*t*_. Black curve: total extinction rate; gray curve: extinctions due to cascade effects. Shaded regions around the average rate curves represent central 95% credibility intervals beta distribution for establishments in A, gamma distribution for extinctions in B]. Panel C: Extinction cascade size distribution. Each bar represents the relative frequency for extinction cascades of the corresponding size; the lightness level is determined by the percentage of extinctions due to cascade effects, i.e., non-competitive interactions. Exemplary values for these percentages are stated explicitly. Panel D: Smoothed time series of consumer establishments (green), extinctions (black), the corresponding subset of non-competitive exclusions (gray), and the loss rate of resources (red). Smoothing was performed with a Blackman window (size 15 for *t* ∈ [0, 500], size 100 for *t* ∈ [9.5, 10] × 10^4^) as described in Ref. 49].

#### Displacement

During the simulation run shown in Fig. 3, the consumer richness saturates after an initial growth phase of the network and fluctuates around finite values of *N*_*t*_ ≈ 125 *< N*_max_, where *N*_max_ (*A* = 0) = 233, cf. (17). This constraining of the richness below its maximal theoretical value corresponds to a balance of successful establishments of new species and extinctions of previous residents. Figure 5 A shows that the average establishment success rates for different values of the consumer richness is slightly decreasing. However, it is larger than 0.5 for all community sizes observed in the simulation, indicating that at least every second newly generated consumer is able to establish for any of the observed community sizes. Hence, the ratio of average extinction to establishment rate must increase with larger resident community sizes *N*. Because extinctions can only occur as a consequence to a new establishment, this is equivalent to the extinction rate conditional to species establishment exceeding one. See Panel B to observe, that this conditional extinction rate (black curve) is relatively close to one for a range of community sizes *N* ∈ [95, 1 50], which indicates that the consumer richness drifts up and down within this region. However, an increase of the average extinction rate around *N* = 160 sets an upper limit to these fluctuations. Here, only a few samples exist for *N >* 160, which causes enlarged 95% credibility intervals (shaded hull of the extinction rate curve).

For the observed extinction events, we may differentiate between two types of extinctions. Firstly, we classify events as competitive exclusion if the extinct consumer is a direct competitor to the newly introduced consumer, i.e., extinct and new consumer share at least one resource. Secondly, there are indirect effects, which may cause extinctions by the altered resource flow pattern at a different network location. For instance, reduced resource flows may result due to extinctions or increased competition for their producers. We call the category of the latter phenomena “cascade effects”. Figure 5 B and D differentiate between these two cases, showing the fraction of extinctions attributed to cascade effects as a lighter gray curves. The total fraction of extinctions attributed to cascade effects in this sense is about 14.8%. It is larger for steps where more than one consumer goes extinct.

The distribution of the extinction cascade sizes is shown in Fig. 5 C. Here, the corresponding bars are shaded by the fraction of extinctions attributable to the direct competition with the new consumer. For comparison, these fractions are annotated for selected bars. Fitting an exponential distribution, we obtain a decay of cascade size frequencies with an exponent of *−*0.27.

### 3.3 Parameter sensitivity

In this section, we assess the effect of a number of selected parameters on the evolution of the CFN. Figure 6 shows various measures for an array of numerical simulation experiments with specific parameter sets. Each row contains information about four different experiments, which implement a gradient for the value of a specific parameter, see legend on the left. All other parameters were fixed to the values stated in the caption of Fig. 3. The reported quantities are averaged over the step interval [5, 10] × 10^4^.

**Figure 6:**
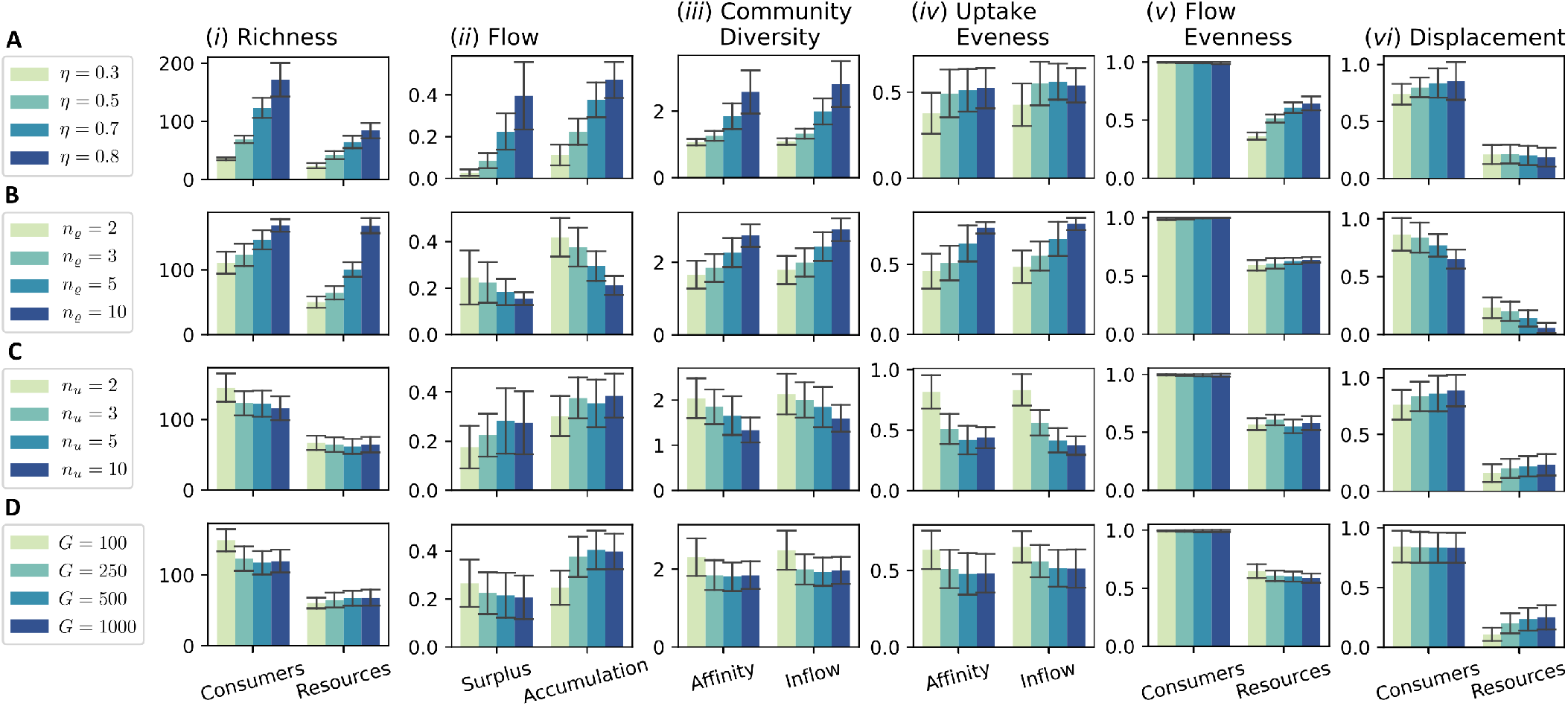
Parameter dependence of macroscopic measures. The rows A-D each refer to the variation of a particular parameter A: recycling fraction *η*; B release diversity *n*_*ϱ*_; C uptake diversity *n*_*u*_; D global resource pool size *G* = # 𝒢]. Different colors indicate different values for the corresponding parameter (see legends). If not indicated otherwise, all parameters are fixed to the values given in Fig. 3. Bar heights in the columns (i)-(vi) show the mean values of different observables in different simulation runs averaged in the step interval [9.5, 10] ×10^4^, errorbars delineate an interval of two standard deviations around the mean. (i): Average consumer and resource richness 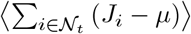, and ⟨*M*_*t*_ ⟩; (ii): Average total surplus flow⟨ ∑ _*i*∈𝒩_*t* (*J*_*i*_ *− µ*) ⟩, and average accumulation flow ⟨*A*_*t*_⟩, cf. (14); (iii) Community diversity based on affinities and on uptake flows, cf. (21); (iv): Uptake evenness ⟨*E*_*t,i*_ ⟩averaged over consumers and time, where 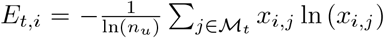 ln (*x*_*i,j*_), with 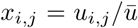 (affinity) or *x*_*i,j*_ = *J*_*j*→*i*_*/J*_*i*_ (inflow); (v) The average ensemble evenness of consumers and resources based on the flows 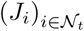, and 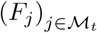, respectively; (vi): Average displacement rates (smoothed over 100 steps as in the second interval of Fig. 5 D) for consumers and resources.

#### A. Variation of the recycling fraction

*η* In the first sequence, cf. Fig. 6 A, we tested four different values for the recycling fraction *η*, ranging from 0.3 to 0.8. Recall that the flow scales with (1 *− η*)^−1^, cf. (13). This is resembled by the observed consumer richness, which increases with *η*, cf. Panel A(i). As a consequence of the higher consumer richness and the corresponding diversity of released resources, the number of different resources increases as well. Less predictable, we also observe a clear increase in the accumulation and as well in the surplus flow, cf. Panel A(ii). The surplus fraction increases more than the consumer richness, which means that, on average, a consumer’s uptake flow has a larger buffering towards the extinction threshold *µ*. This is not clearly reflected by any decrease in the average extinction rate, which instead show a slight increase, cf. Panel A(vi). On average, the diversity of the community and also the average evenness of assimilated resources per consumer increase with rising *η* cf. A(iii) and A(iv)]. This seems to be a consequence of the increasing number of consumers dwelling on secondary, non-supplied substrates, since we observe the same pattern of consumer richness being correlated to community diversity and uptake evenness across all sequences A-D. An associated effect is the increase in the resource flow evenness, resulting from a higher production flow through non-supplied resource pools, see A(v).

#### B. Variation of the release diversity

*n*_*ϱ*_ Similar as for the sequence A of varying recycling fraction *η*, we find an increase in consumer and resource richness for higher values of *n*_*ϱ*_ in sequence B. In contrast to A this increase is more pronounced in the resource richness. The latter seems intuitive: as each consumer produces a higher number of resources, the total pool of resources grows. That this leads to a higher consumer richness, seems to be a consequence of a specific flow pattern, different to the one in case A. While the consumer richness increases [see B(i)], the flow per consumer decreases and likewise does the fraction of flow into accumulating resources [see B(ii)]. Thus, a higher percentage of the flow is assimilated and more equally distributed among consumers for higher values of *n*_*ϱ*_. The reason for this more efficient exploitation of the system’s resources seems to be the increased stability resulting from the fact that produced resources tend to be formed by more different producers. Therefore, a single species’ extinction is less severely affecting the environmental conditions, allowing the evolutionary adaptation to keep the pace of environmental change. This stability is also reflected in the significantly reduced extinction rates for higher *n*_*ϱ*_ [see B(vi)] and a decreased average evolutionary potential (by a factor of ≈ 2.34, not shown). This finding stands in contrast to the classic argument of May [50] relating an increased diversity to a decreased stability, but is in accord with recent findings in food web models where the size of trophic cascades was reduced with the complexity of the food web [51].

#### C. Variation of the uptake diversity

*n*_*u*_ Regarding sequence C of Fig. 6, one might expect, that an increase of the uptake diversity *n*_*u*_ would have similarly stabilizing consequences as an increased release diversity *n*_*ϱ*_ (see case B), since a consumer could now be able to hedge its necessities across a higher diversity of compounds.

However, in contrast to case B, the consumer richness decreases, while the average extinction rate slightly increases. A difference between the parameters *n*_*Q*_ and *n*_*u*_ is that *n*_*u*_ controls the dimension of the parameter space’s portion most directly relevant to the consumer fitness, i.e., the value of its uptake flow *J*_*i*_. Thus, we conjecture that this is a potential source for the increased instability of systems with increased uptake diversity *n*_*u*_. Since the exploration of the affinity parameter space is more difficult, this decreases the pace of evolutionary adaptation. At the same time, the impact on environmental conditions per extinction is, insofar it is controlled by *n*_*Q*_, the same in all considered scenarios of sequence C. Indeed, the mean value of the average evolutionary potential (19) is maximal for the case *n*_*u*_ = 10 among all considered simulations and about 5-fold as high as the value in the reference case *n*_*u*_ = 3 displayed in Fig. 3.

#### D. Variation of the global resource pool size

*G* In Fig 6 D, we have varied the size *G* = #𝒢 of the global resource pool. For any given state of the CFN, an increase of the global resource pool would lead to a higher expected number *M*_*t*+1_ of resources after the introduction of a new consumer. Assuming a uniform probability of resources to be chosen upon mutation, the probability for a new resource to be chosen as a release product of the new consumer equals 1*/G*. Accordingly, the probability that this choice falls into the part of previously absent resources is (*G − M*_*t*_) */G*. This suggests that the average resource richness would increase with *G*. On the contrary, we observe no significant dependence of the average resource richness with rising *G*, see D(i). The reason why the naive local analysis offered before fails is due to network effects that must be taken into account. In fact, the emergence of a new metabolite creates an additional flow into unused, accumulating resource pools, see D(ii). This accumulation remains until a consumer evolves, which is capable to utilize the new resource. As new metabolites emerge more frequently, they also vanish more frequently in the statistically stationary state cf. D(vi)]. This elevated turnover provides a less stable environment. Given less time for adaption, this leads to a less efficient exploitation. In effect, the decreased number of producers compensates the effect of increased resource diversification rate at establishment events and the resulting resource richness remains approximately constant.

### 3.4 Cluster dynamics

In our framework, consumers may be more or less similar, or even identical, to each other, if compared on the level of their metabolic characteristics. To characterize and compare the structure within different communities and its change over time, we pursue an approach based on ancestral relations and similarity clustering of consumers. A cluster in our context may be considered to correspond to a taxonomic unit consisting of different variants or closely related strains of a common taxon.

Based on such an approach, we obtain a representation of evolutionary change as shown in Fig. 7 for different parameter sets. Every node in the diagram represents a cluster of similar consumers at a corresponding step in evolutionary time. A link between two clusters at subsequent time points is postulated if an ancestral relationship between these clusters exists. We start with an initial hierarchical clustering of a community at a given initial step *t*_0_. The dendrograms on the left of each timeline show this initial clustering, which does not take into account any ancestral information. The subsequent development of clusters through time is defined as follows. If a new consumer establishes itself in the community, it is added to the cluster its direct ancestor belongs to. However, if the diameter of the extended cluster exceeds a specified threshold *?*, it is split into two, assigning a slightly altered color to the part containing the new consumer. For the initial, as well as for the separative, clustering, we use a hierarchical approach 53] based on the pairwise “genotypic” dissimilarity of two consumers defined as

**Figure 7:**
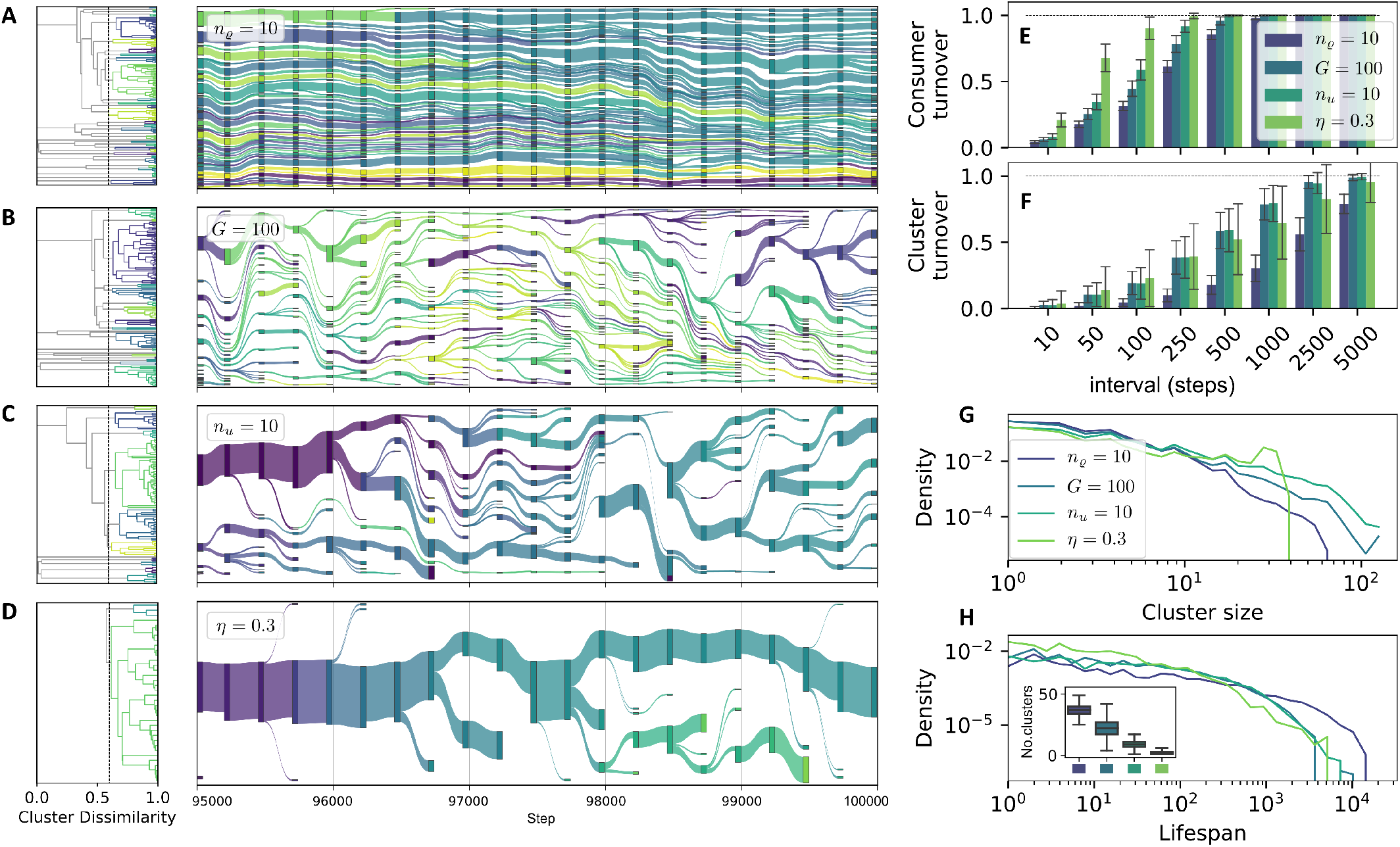
Dynamics of similarity clusters of consumers. A-D: alluvial diagrams of cluster lineages. The link weights between two ancestrally related clusters at steps *t* and *t* + 1 are computed proportional to the relative size of the cluster at step *t* + 1. Dendrograms to the left show the initial clustering of the community at step *t* = 95000 used as a basis for the remaining evolutionary tree. We use *ϑ*= 0.4, *w*_*u*_ = 1.0, *w*_*ϱ*_ = 0.2. Other parameters as in Fig. 3 or indicated in the legend. Visualization with plotly [52]. Panels E and F: Average turnover statistics comparing communities 𝒩_*t*_ and 𝒩_*t*+*I*_ (in E) and present clusters (in F) for different step intervals *I*, cf. Eq. (24), and *t* ∈ [5, 10] × 10^4^. Error bars show the standard deviation over all turnover values measured at the corresponding intervals. Panel G shows the densities of the logarithmic cluster sizes for the different cases. Panel H displays the densities of logarithmic cluster lifespans with and inset of the observed number of cluster (the boxplot indicates the 5%, 25%, 50%, 75%, and 95% quantiles of the distribution).

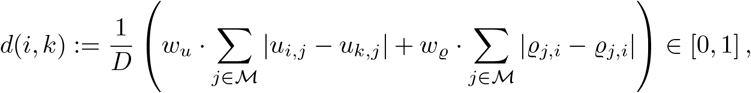

for any pair (*i, k*) of consumers, where 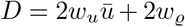. To generate the cluster hierarchy cf. Fig. 7(e)], we define a dissimilarity between two clusters *c*_1_ and *c*_2_ as

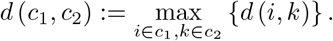

Figure 7 A-D shows four representative cases of community dynamics with different features for the step interval [9.5, 10] × 10^4^. The depicted cases correspond to variations in cluster stability and diversity. Panels E-H show several statistics for the system’s evolution in cases A-D. Panels E and F show the turnover of consumers and clusters against time, where the turnover between two sets *A* and *B* which contain consumers in E and clusters in F] is calculated as [54]

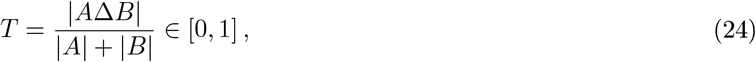

where *A*Δ*B* denotes the symmetric difference of *A* and *B*. Panels G and H show the densities of the logarithms of the cluster sizes (in G) and of the cluster lifespans (in H).

Panel A of Fig. 7 displays a case of high diversity and high stability, corresponding to the parameter variation of high release diversity *n*_*ϱ*_ = 10, which is also reported in Fig. 6 as part of the sequence B. For this case, the turnover on consumer level as well as on cluster level is the lowest among all depicted cases in Fig. 7, cf. Panels E and F. The average lifespan of clusters is the highest, cf. H. This value is computed as the number of steps between a clusters first appearance and its disappearance when its last member goes extinct. Due to their relatively high number, the individual clusters are smaller than in most other cases, cf. G and H.

Cases B and C correspond to the parameter variations of small global resource pool size *G* = 100 and of high uptake diversity *n*_*u*_ = 10. Both cases exhibit intermediate turnover rates on consumer and cluster level, which can be associated with the intermediate consumer displacement rates reported in Fig. 6 C(vi) and B(vi) for both cases. Interestingly, we see a higher consumer turnover for *n*_*u*_ = 10, while the cluster turnover is approximately equal for both cases. At first glance, one might think that the decreased resource pool turnover for the case *G* = 100 cf. Fig. 6 D(vi)] could stabilize the environmental conditions, allowing for more gradual shift in the community composition. Such gradual shifts would correspond to more constant clusters, because a cluster could gradually follow the environmental shift without splitting up. On the contrary to these considerations, we find a relative decrease of the cluster turnover for *n*_*u*_ = 10, i.e., the ratio of cluster turnover to consumer turnover is smaller for this parameter variation. Note that the case *G* = 100 exhibits a higher richness and diversity see Fig. 6 D(i) and D(iii)], and on average more clusters cf. Panels B and G], which might be responsible for relative increase in the cluster turnover.

An even smaller ratio of cluster to consumer turnover cf. Panels E and F] is displayed by the parameter variation *η* = 0.3, where we find the highest consumer turnover rates, while the cluster turnover, especially for longer time intervals, is even lower than for the cases B and C. The distinguishing characteristic of the cluster structure for the case *η* = 0.3 is the concentration of almost all consumers in a single, large and stable cluster, which leaves its trace in the cluster size distribution as a bump around a value of 120, see Panel G. This low cluster diversity emerges from the low diversity of resources and consumers for low recycling fractions *η* - cf. Fig. 6 A and the corresponding discussion in Sec. 3.3. Due to the scarcity of different niches, only a few branchings from the main cluster of supply consumers take place. Smaller clusters disappear in the majority of cases, indicating a gradual evolution rather than large innovative steps.

## 4 Discussion

### Model Assumptions and Limitations

Several idealizations or simplifications were assumed in the derivation of the model. Perhaps, the crudest simplification may seem to be the limitation of the number of different organic compounds to a maximal global resource pool size *G*, which lies orders of magnitude below the number observed in natural systems. For instance, the oceanic DOM was estimated to be composed of more than 100, 000 different compounds 55]. However, the effect of larger resource pool sizes in our model is that the probability for each newly evolved release substrate to be already present in the system tends to zero. Apparently, this limit is already well approximated by a resource pool of size *G* = 500 in our setup, which does not lead to significantly distinct behavior compared to *G* = 1000, cf. Fig. 4 D. The size, where an increase of *G* does not lead to significantly different evolutionary dynamics, may increase with the number of consumers and the diversity *n*_*Q*_ of released resources per consumer species, but whether this could lead to qualitative differences in parameter dependencies is questionable.

As another simplification, we assumed no particular properties for any of the resources aside of the designation of a supply point. No preferential attachment or evolutionary drive towards the most abundant resources was assumed beyond that. Further, each resource had the same nutritional value for each consumer. Indeed, neither the assimilation efficiency was varied, i.e., *η*_*i,j*_ ≡ *η*, nor was the total affinity *ū* assumed to depend on the consumed resources, which might correspond to different maximal uptake rates for the different resources. Moreover, we neglected any abiotic degradation of substrates. This does not imply qualitative differences beyond neglected variations in uptake efficiencies for resources, which are consumed. With some modifications, a corresponding term can be integrated in the formalism of Sec. 2.1, resulting in a slightly modified expression for the transformation matrix T. However, if the resource concentrations were to be determined, released, but unconsumed resources would accumulate to infinity, when degradation is disregarded. This is not important for the part of interest in this paper, though, since this omission does not create any singularities in the network flow, and resource concentrations are not assumed to affect the development of new consumption capabilities.

The most important characteristic displayed by the model, i.e., the ongoing co-evolution, is not affected by the above simplifications, because the fundamental mechanism, driving the continuous turnover of the presented CFN, does not rely on any of these simplifications. This mechanism is the appearance and disappearance of produced metabolites, which is subject to an undirected evolutionary drift, that constantly alters the environmental restrictions of its beneficiaries, i.e., consumers living on that products. The drift of release configuration is undirected, because the producer’s own fitness, that is its associated inflow *J*_*i*_, is largely independent of the of the exact configuration of its products.

The evolutionary dynamics displayed in our model do not stably reproduce all types of resource-mediated ecological relationships. For instance, there is no stabilizing mechanism for reciprocal cross-feeding, which is observed in microbial communities. To see that, consider the simple scenario of a pair of consumers *i* and *k*, consuming each other’s products. In this situation any consumer *i*_*′*_ with affinities leading to *J*_*i′*_ *> J*_*i*_ has the potential to displace consumer *i*, even if it does not produce any of the resources consumed by *k*. This is a variant of the tragedy of the commons [56], since each produced resource in our model is in principle accessible to each consumer. The development of stable reciprocal cross-feeding in natural microbial communities is believed to be a stepwise process [44, 57]. The first step is an accidental bidirectional exploitation of metabolic waste products as it also may arise in our model. However, the stabilization and optimization of the syntrophic relationship involves processes beyond the scope of our model. For instance, a spatial association may stabilize a pair of cooperative strains as has been demonstrated recently[36, 58]. Under which circumstances such a stabilization could counteract the perpetual changes exhibited by our model is not obvious and could be a subject for further studies.

A distinctive feature of our model are the inelastic, quantized affinities for the different consumers. We do not explicitly model a variable population density for each consumer, but arbitrate its presence or absence by a consultation of Eq. (12). In a simple dynamical growth model, with mortality *µ*, the difference *J*_*i*_ *− µ* could describe the instantaneous, per capita growth rate for a population of the corresponding consumer. Therefore, Eq. (12) would result in a negative growth rate in that framework. However, since consumers tend to produce similar descendants, a cluster of similar consumers quickly populates any available resource. Translating a cluster into a framework of variable population densities, it corresponds to a population with an internal variability. Although the quantized representation of population densities is a simplification with respect to continuous representations, it reflects the intraspecific variation inherent to the concept of a taxonomic unit [59].

### Resource Diversification and Community Turnover

Our CFN self-organizes towards a high level of diversity in both substrates and microorganisms, even after starting with a single microorganism in the in silico experiment. Diversification occurs in two aspects in our model, which are both in line with empirical evidence: microorganisms transform and release compounds that can serve as substrates for other microorganisms, i.e., they diversify compounds, and a diverse set of compounds in turn can support a diverse microbial community. Indeed, the exometabolome of heterotrophic microorganisms shows a remarkable diversity, even when grown on a single substrate [60-62], providing potential new niches for other microorganisms. In this context, the role of enzyme promiscuity has been highlighted as a possible mechanism to potentiate the number of transformed resources and has been suggested to contribute to the huge diversity of organic compounds dissolved in seawater[9]. This ability of enzymes to produce new, promiscuous products and to work on substrates besides their canonical substrate spectrum, may trigger the evolutionary development of new metabolic pathways [63]. Evolutionary diversification of substrate associations has been observed in a laboratory experiment, in which *E. coli* was grown on glucose as a single substrate under static environmental conditions. After more than 400 generations, strains evolved that did not primarily depend on glucose anymore but rather on metabolites exuded from *E. coli* [20, 21]. Bacteria seem to evolve even more rapidly when interacting with other species compared to evolution of isolated strains [64]. Our results corroborate the hypothesis that this high diversity of substrates and microorganisms is self-sustaining and a direct result of cross-feeding interactions subject to evolutionary dynamics.

While the high level of diversity remains fairly constant, the actual microbial and molecular community is constantly turned over. With the constant community shift by the appearance or vanishing of new substrates, the fitness of individual microorganisms varies with each establishment of a new species. Experimentally, microbial selection and community assembly has mainly been studied with regard to changing environmental conditions. Only few studies specifically targeted microbial community turnover at stable conditions. In line with our results, a continuous turnover under static conditions has been directly or indirectly suggested: Danczak et al. 65] found an uncoupling between environment and the development of microbial communities in unperturbed aquifers, leading them to hypothesize that unperturbed environments house dynamic communities due to external and internal forces. Similarly, Konopka, Lindemann, and Fredrickson 66] concluded that probably no environmental community truly reaches steady-state but exists in a constant flux, which is in line with a metaanalysis by Shade et al. [67]. Our results suggest that an ongoing evolution of syntrophic or commensal interactions is a potential internal mechanism, leading to the observed constant turnover of microbial community composition.

### Microbial Community Structure

Despite the constant turnover, the community composition in our simulation experiments shows the emergence of specialist and generalist consumers. Here, a microorganism in the network is considered a generalist if the uptake is distributed evenly among different substrates, whereas specialists mainly rely on a single resource. Although our model is deliberately set in a static environment, we observe both strategies non-randomly distributed in the network. Specialists occur in network locations around a less diverse but highly concentrated inflow, i.e., at the constantly externally supplied resource, cf. Fig. 4 B. On the other hand, resources that constitute one ingredient in a diet of a generalist usually exhibit significantly lower inflows, and fluctuate stronger when new species establish. These fluctuations are results of evolutionary drifts of the released resources, which do not directly, and much more weakly, affect the realized fitness of a consumer. The slow, constrained evolution of affinities, which optimizes a sub-community on a persistently supplied resource, is accompanied by an unconstrained, rather rapidly drifting combination of products that are formed from the metabolization of the more persistent resource. In line with our simulations, the experimental evolution of a bacterium (Pse*u*domonas f*iu*orescens) showed that in complex environments with many substrates, neither narrow specialists nor complete generalists evolve but rather overlapping imperfect generalists, which have adapted not to all but a certain range of substrates [68]. Several examples of heterogeneous microbial communities seem to show strategies of evolving to optimize bet-hedging for substrates [69-71]. Observations from stability patterns in wastewater treatment plants show similar dynamics to the emergence of characteristic patterns resulting from differences in the resource flows in our model [72]. Our results indicate that these patterns may already emerge from the basic principles of diversification and evolution that our idealized CFN model is set on. A more decisive exploration of the relation between fluctuation of abiotic factors and evolving patterns of resource specialization in CFNs represents an interesting subject to further research.

Additionally, microbial networks often show a clustered topology, e.g., in soils [73] or oceans [74], reminiscent to the clusters observed in our and other author’s models. For instance, Goyal and Maslov[[37] reported that their model is capable of reproducing a structure of a core community, which contains species occurring with high abundance, combined with peripheral, less abundant, species. For a range of parameter values, our model reproduces such a distribution corresponding to unequal cluster sizes, which follow the magnitude of flow associated with the utilized resource pools (most apparent in Fig. 7 D). In addition to the results presented in [37], we found more diverse network structures depending on the system parameters. According to our results, the diversity of released resources *n*_*ϱ*_, and the fraction of recycled flow *η* are the most significant factors for the structure of the resulting communities.

It is worth noting that our - as well as Goyal and Maslov’s - results do not rely on any inherent biochemical nature of the different resources. Although the magnitude and diversity of the externally supplied resource flow influences the complexity of a resident community, in our model a complex community emerges from a single, basal resource that is externally supplied. One may ask: how much of the structure of natural communities is emergent, and how much is an artifact of an underlying restriction of resource transformations or, more general, of the constraints of the chemical universe The molecular space possesses a wealth of molecular structures, and microbial metabolism is subject to chemical constraints such as elemental stoichiometry of uptake and release, energetic gains during heterotrophic metabolic transformations or preferences of macromolecules sich as sugars, lipids or amino acids [43]. These constraints define possible combinations of uptake and release configurations [16, 40]. As our results show that a structured CFN may develop without such assumptions, we conclude that the degree of correlation between resource availability with respect to the abovementioned constraints, and the community of microbial consumers is not necessarily high. Thus, a quantification of the emergent fraction in community composition represents an important matter of further research.

In summary, we studied a model for the evolutionary change of CFNs and presented a numerical analysis of the influence of several parameters on emerging structural properties. Our model provides a framework to study natural evolutionary and radiation processes in microbial ecosystems and shows that, when combined, these do not lead to a stable microbial community but rather to a persistent co-evolution and turnover, even when environmental conditions remain static. We showed that, starting from one microbial consumer feeding on a single substrate, CFNs can self-organize into systems with diverse, coexisting microorganisms producing and feeding on numerous substrates. The diversity of the system is self-sustaining because new microbes produce new substrates, which in turn allow for the establishments of new microbes. While the community is constantly turned over, the diversity of compounds and microbes eventually saturates, where the asymptotic diversity of resources depends strongly on the microorganism’s capability to diversify substrates.

